# The Spatial Organization of Bacterial Transcriptional Regulatory Networks

**DOI:** 10.1101/2022.07.19.500698

**Authors:** Liu Tian, Tong Liu, Kang-Jian Hua, Xiao-Pan Hu, Bin-Guang Ma

## Abstract

Transcriptional regulatory network (TRN) is the central pivot of a prokaryotic organism to receive, process and respond to internal and external environmental information. However, little is known about its spatial organization so far. In recent years, chromatin interaction data of bacteria such as *Escherichia coli* and *Bacillus subtilis* have been published, making it possible to study the spatial organization of bacterial transcriptional regulatory networks. By combining TRNs and chromatin interaction data of *E. coli* and *B. subtilis*, we explored the spatial organization characteristics of bacterial TRNs in many aspects such as regulation directions (positive and negative), central nodes (hubs, bottlenecks), hierarchical levels (top, middle, bottom) and network motifs (feed-forward loops and single input modules) of the TRNs and found that the bacterial TRNs have a variety of stable spatial organization features under different physiological conditions which may be closely related with basic life activities. Our findings provided new insights into the connection between transcriptional regulation and the spatial organization of chromosome in bacteria, and might serve as a foundation for spatial-distance-based gene circuit design in synthetic biology.

## Introduction

A transcriptional regulatory network (TRN) is composed of the regulatory relationships between transcription factors (TF) and their target genes which serves as the information processing hub to respond to intracellular and environmental signals [1–3]. The regulatory relationships in TRN could be positive (activation) or negative (repression) or both (conditionally activation or repression) and the whole TRN is usually scale-free with small numbers of highly connected nodes [4]. Since TRN is a directed graph, it is usually organized in a hierarchical structure and genes of different levels in the hierarchy usually bear different functions; a hierarchical structure makes TRN more robust [5, 6]. There are building blocks in TRN called network motifs which are connected subgraphs that appear more frequently in real networks than in the corresponding randomized networks. Typical network motifs in TRN are negative auto-regulation, feed-forward loop (FFL) and single input module (SIM). The specific topologies of these network motifs give them unique dynamic functions; for example, FFLs have the function of filtering noisy signals and speeding up responses, while SIMs can switch ON/OFF multiple target genes in a temporal sequential order, often acting as a timer during biological development [7]. In TRN, various network motifs are intertwined to produce rich kinetic functions, which jointly regulate the growth and development of an organism.

In previous studies of kinetic modeling, the distribution of genes in space and the time required for transcription factors to search for target genes were often ignored [8]. In recent years, with the development of single-molecule fluorescence tracer technology, it has been found that transcription factors search for their target genes much longer than expected, and the time required for a single lacI protein to find its target gene takes 3-5 minutes [9], while a single dCas9 protein finding its target gene takes even 5 hours [10]. This is because the higher the specificity of a transcription factor to its binding site, the lower the search speed, and a compromise is needed between these two. In addition, the spatial distance between a transcription factor and a target gene significantly affects the rate and reliability of transcriptional regulation in bacterial cells, and shorter search time leads to higher regulation efficiency [11]. When a transcription factor is far away from its target gene, the transcription factor may be unable to find the target gene [12]. Moreover, functionally related genes such as the genes whose products participate in protein-protein interactions or in the same metabolic pathways were found spatially clustered in cell [13, 14]. The spatial distance between genes also affects the dynamics of gene network. Van *et al*. studied the dynamics of three-node Repressilator and found that different distribution patterns of the three genes in space will affect whether the oscillation behavior can appear and stably exist [15]. It has been acknowledged that the chromosome architecture affects the basic biological processes of bacteria such as replication, transcription and gene regulation [16, 17]. On the other hand, these processes also in turn affect the chromosome architecture through evolutionary pressure [18–20].

In recent years, chromatin interaction data of bacteria such as *Escherichia coli* [21, 22] and *Bacillus subtilis* [23–25] have been published, which makes it possible to study the spatial organization of transcriptional regulation on the scale of the entire transcriptional regulatory network. *E. coli* and *B. subtilis* are model strains of Gram-negative and Gram-positive bacteria, respectively, with clear genetic background and relatively complete transcriptional regulation data. At the same time, the 3D architecture of chromosomes, the spatial distribution of RNA polymerase and ribosomes, and the growth and replication processes are quite different for these two bacterial species [16, 26]. In this study, *E. coli* and *B. subtilis* were used as the representative species of bacteria and their TRNs were constructed and combined with the chromatin interaction data under multiple culture conditions. The spatial organization of TRN was explored in terms of different TRN features such as regulatory direction (positive and negative), central nodes, network hierarchy and network motifs. It was found that there are stable organizational features of TRN under various conditions for the two bacterial species. The results provided new insights into the organization principles of TRN in 3D physical space and might guide spatial-distance-based gene circuit design in synthetic biology.

## Results

### Chromatin interaction within TRN

To explore the spatial organization of bacterial TRN, the chromatin interaction data and TRNs for the two model organisms *Escherichia coli* and *Bacillus subtilis* were utilized. For *E. coli*, the chromatin interaction data were measured at 5 culture conditions [22] and denoted as E-LB30, E-LB37, E-MM22, E-MM30, E-sMM30, correspondingly; for *B. subtilis*, the chromatin interaction data were measured at 3 culture conditions [23] and denoted as B-LB, B-MM, B-Rif, correspondingly; see **Methods** for details. The TRNs were constructed based on the data from RegulonDB (version 9.0) [27] for *E. coli* and from SubtiWiki2.0 [28] for *B. subtilis* (**Figure S1**, see **Methods** for details). First, the chromatin interaction frequencies within TRN (namely, between the regulators and their target genes in the two bacterial TRNs) (denoted as TRN) were calculated and then compared with the chromatin interaction frequencies among all the gene pairs in the whole genome (denoted as All). As shown in **Figure 1 and Table S1**, TRN is significantly higher than All, meaning that the genes with regulation relationships tend to be closer to each other in the 3D space. This result extends our previous finding that genes in the same regulon or pathway of *E. coli* tend to minimize their spatial distances [14], to the genes in the TRNs of both Gram-negative (*E. coli*) and Gram-positive (*B. subtilis*) bacterial species.

**Figure 1.**
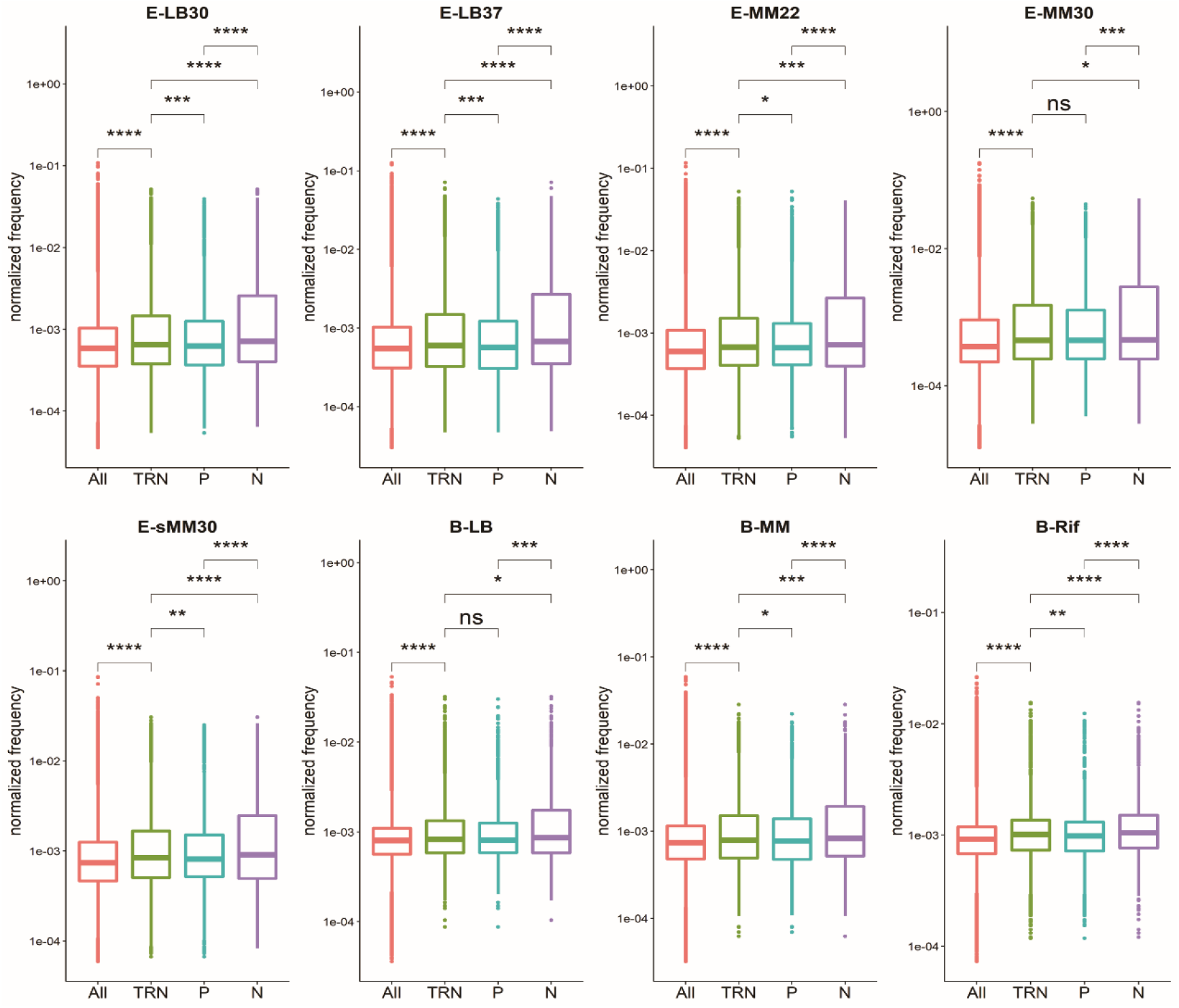
Comparison of chromatin interaction frequency in global TRN. Under almost all culture conditions, the chromatin interaction frequency within TRN (denoted as TRN) is significantly higher than that of all the gene pairs in the whole genome (denoted as All); the interaction frequency of positive regulation (denoted as P) is significantly lower than TRN; the interaction frequency of negative regulation (denoted as N) is significantly higher than TRN.

We compared the chromatin interaction frequencies of positive and negative regulations with TRN in the two bacterial TRNs (**Figure 1 and Table S1**). We found that the interaction frequencies of positive regulation in almost all culture conditions were significantly lower than the interaction frequency within TRN, while the interaction frequencies of negative regulation were significantly higher than the interaction frequency within TRN. It suggests that negative regulation (repression) prefers shorter spatial distance while positive regulation (activation) is just the opposite.

### Chromatin interaction of central nodes in TRN

In TRN, different nodes are of different importance and centrality can measure the significance of a node’s impact on the network topology or biological function. We calculated the in-degree, out-degree, betweenness and closeness centrality values for each node and selected the nodes with the top 20% highest centrality values as the central nodes (42 for *E. coli* and 38 for *B. subtilis*), denoted as In-Hubs, Out-Hubs, Bottlenecks and Centers, respectively. We examined the chromatin interaction frequencies intra the central nodes and between the central nodes and other connected nodes in TRN. Compared with the chromatin interaction frequency of TRN (**Figure S2**), we found that the chromatin interaction frequencies intra In-Hubs, Out-Hubs and Bottlenecks are usually higher than that of the TRN, while the chromatin interaction frequencies between the Out-Hubs, Bottlenecks and their corresponding connected partners are usually lower than that of the TRN. The frequency of chromatin interaction that centers participate in seldom has significant difference with that of TRN for most of the culture conditions of the two species.

The TFs in Out-Hubs and Bottlenecks need to regulate a large number of genes and usually have a high expression level. For instance, sigma factors are the essential TFs for transcription initiation [29] and involve in more than a quarter of regulation edges. Most of the sigma factors (6/7 in *E. coli* and 13/20 in *B. subtilis*) are attributed as central nodes. Therefore, shorter spatial distances between central nodes imply clustering of these genes (TFs) in space, which may benefit from the so-called transcription factories [30] in bacteria to fulfill cost-effective transcription. On the other hand, a huge number of genes are regulated by central nodes, which means these genes are widely distributed in the genome, resulting in relatively lower (than TRN) interaction frequencies between the central nodes and their connected partners.

### Chromatin interaction in the hierarchy of TRN

According to the network topology, TRNs can be divided into four layers: Top, Middle, Bottom and Target (see **Methods**). Except for the Target layer, the genes in the other three layers are transcription factors. In the Top, Middle and Bottom layers (**Figure S3**), there are 8, 75, 131 genes for *E. coli* and 6, 38, 148 genes for *B. subtilis*, respectively, and the two bacterial TRNs show a pyramidal hierarchy. We examined the regulatory relationships within and between the layers and found that the majority of the Target layer genes in *E. coli* and *B. subtilis* are directly regulated by the Middle layer genes and that the regulations intra the Middle layer are also dense. In contrast, the Bottom layer, even if the number of genes in this layer is more than the Middle layer, has less intra regulations. There are many self-regulating edges (76/89 in *E. coli* and 76/86 in *B. subtilis*) in the Bottom layer and much fewer self-regulating edges (53/335 in *E. coli* and 24/127 in *B. subtilis*) in the Middle layer. It is evident that the Middle layer is the information processing and transmission hub which can make complex decisions. As a mediator level, the Bottom layer plays a cascading role to enrich the regulation forms of Target genes.

The chromatin interaction frequencies within and between hierarchical levels are compared to the chromatin interaction frequency of TRN (**Figure 2, Table S2**). Interestingly, the spatial organization of the Middle-Bottom-Target hierarchical structure shows a high degree of stability in all culture conditions of *E. coli* and *B. subtilis*: the chromatin interaction frequencies intra the Middle layer (Middle-Middle) and intra the Bottom layer (Bottom-Bottom) and between the Bottom-Target layers are significantly higher than that of the TRN; the chromatin interaction frequencies between Middle-Bottom layers and between Middle-Target layers are significantly lower than that of the TRN; others are of no significant difference compared with TRN.

**Figure 2.**
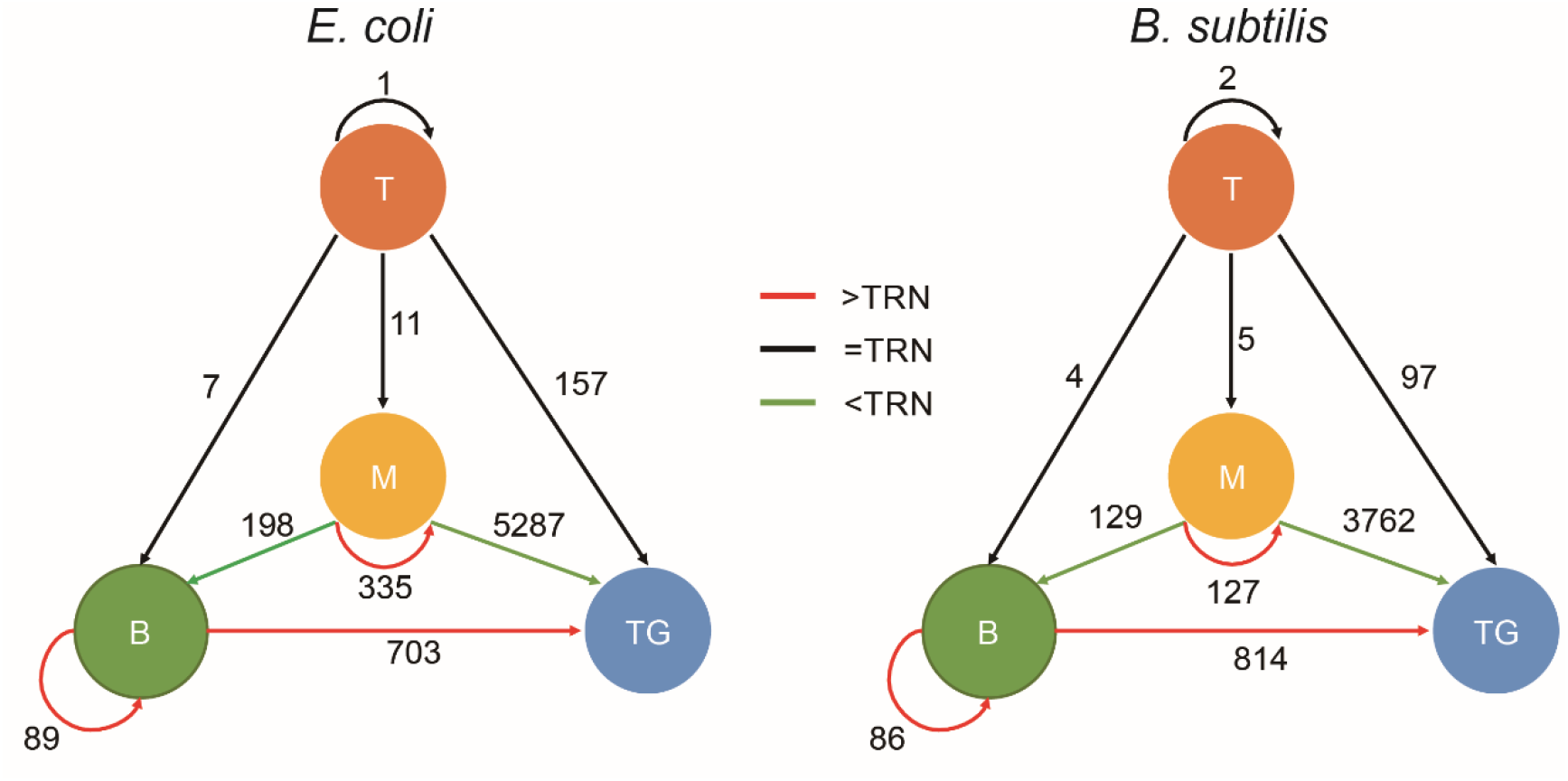
The spatial organization of the TRN hierarchy. The four spheres T, M, B, and TG represent Top, Middle, Bottom and Target layers in the hierarchy, respectively. The numbers on the edges represent the numbers of regulatory relationships within or between layers. The colour of edge represents the comparison result of the chromatin interaction frequency of the edge with the TRN. Significance level: *p* < 0.05.

### Chromatin interaction in the network motifs of TRN

Network motifs are subgraphs with specific dynamic function that are significantly enriched in real networks than in random networks [7]. A total of 4599 FFLs and 30 SIMs were found in the *E. coli* TRN; 2066 FFLs and 58 SIMs were found in the *B. subtilis* TRN (see **Methods**). Only the FFLs with chromatin interaction frequencies on all the three regulation edges and the SIMs with chromatin interaction frequencies on at least two regulation edges are kept for analysis. The chromatin interaction frequencies averaged over all the edges in FFLs and SIMs are calculated and compared with the chromatin interaction frequency of the TRN. Under all culture conditions of *E. coli*, the chromatin interaction frequency within FFLs is significantly higher than that of TRN, while the chromatin interaction frequency within SIMs is of no significant difference from that of the TRN (**Figure 3, Table S3**); under all culture conditions of *B. subtilis*, the chromatin interaction frequencies within FFLs and SIMs were significantly higher than that of TRN. FFLs and SIMs are network subgraphs that perform specific functions and the stronger interaction among their nodes (relative to other genes in the TRN) indicates closer spatial distances among them, which means that the spatial distance is indeed a mechanism to ensure the performance of gene circuit function [15].

**Figure 3.**
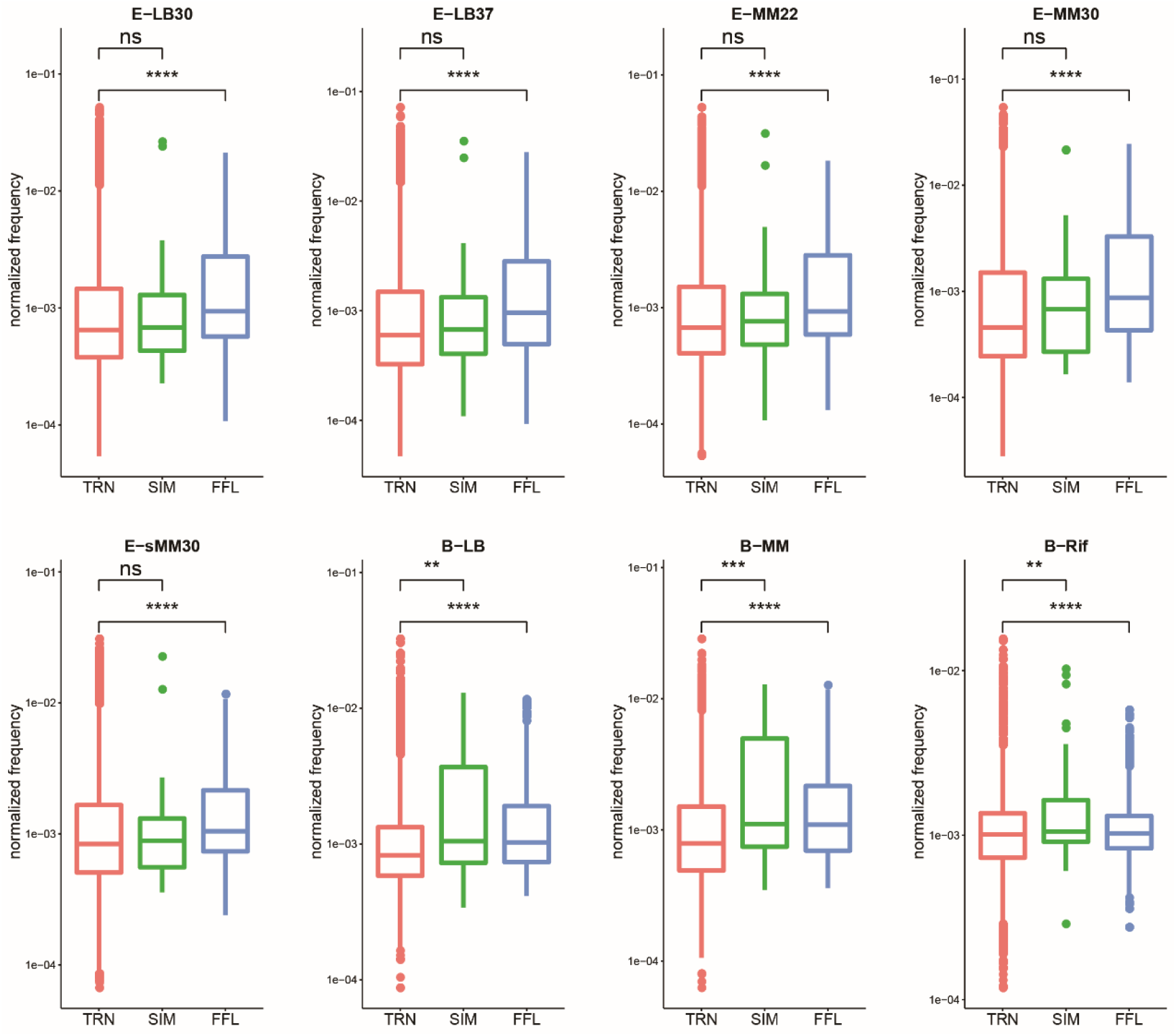
Comparison of chromatin interaction frequencies in network motifs with TRN. Under the five culture conditions of *E. coli*, the chromatin interaction frequency of FFL is significantly higher than that of TRN and the interaction frequency of SIM is of no significant difference from TRN. In *B. subtilis*, the chromatin interaction frequencies of both kinds of network motifs are significantly higher than that of TRN.

If the chromatin interaction frequencies on the three edges in FFLs were considered separately, we found that the chromatin interaction frequencies of the XY and XZ edges are significantly lower than that of the TRN, while that of the YZ edge is significantly higher than that of the TRN (**Figure 4**, **Table S4**), which means that the gene X is relatively farther away from the genes Y and Z and the spatial distance between the genes Y and Z is closer. This mode is highly stable and kept for almost all the physiological conditions of *E. coli* and *B. subtilis*.

**Figure 4.**
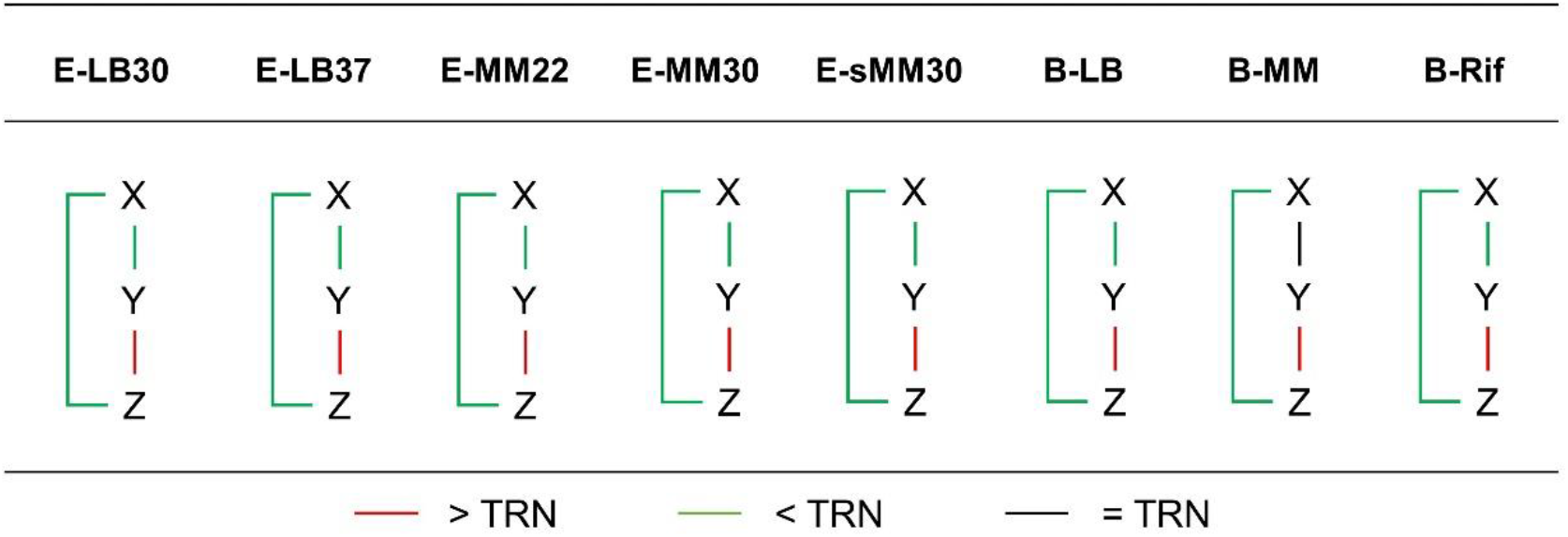
Comparison of chromatin interaction frequencies of FFL edges with TRN. In almost all cases, the chromatin interaction frequency between X and Y/Z in the feed-forward loop is significantly lower than TRN, while the interaction frequency between Y and Z is significantly higher than TRN. Significance level: *p* < 0.05.

By definition, a SIM has one regulator and at least two target genes and can turn ON/OFF the expression of the target genes in a sequential manner owing to the temporal change of the concentration (expression level) of the regulator. As stated above, the chromatin interaction frequency within SIMs (averaged on all edges) is of no significant difference from that of the TRN, which indicates that SIMs may have no special requirement for the absolute spatial distance between a regulator and its target genes. Nonetheless, it was found that the regulation edges within a SIM have similar chromatin interaction frequencies. The standard deviation (SD) of chromatin interaction frequencies on the edges within a SIM was calculated and compared with that of TRN. The statistical results are shown in **Figure 5**. Under all culture conditions of the two bacteria, the SD of chromatin interaction frequency within SIMs is less than that of the TRN. These results show that the chromatin interaction frequency within a SIM is of higher uniformity, which indicates that a regulator in a SIM has similar spatial distances to its target genes and the target genes encounter similar concentration of the regulator at the same time point and the order of state (ON/OFF) switching of target genes is merely determined by the binding affinity between the regulator and the targets.

**Figure 5.**
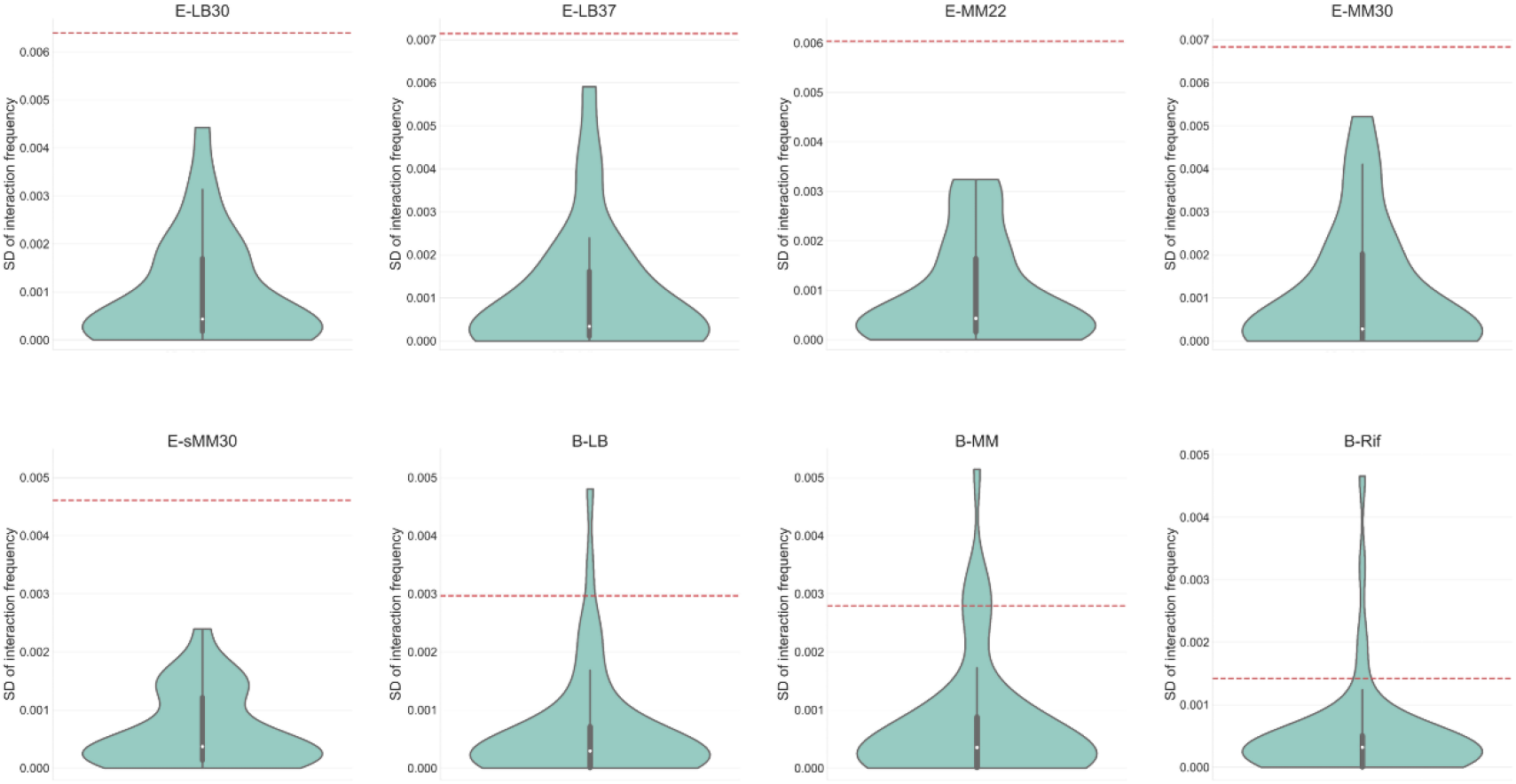
The distribution of standard deviation (SD) of chromatin interaction frequency within SIMs (violin plot) and its comparison with that of TRN (red dashed line). The SD of chromatin interaction frequency within SIMs is apparently lower than that of TRN, indicating lower dispersion and higher uniformity of chromatin interaction within SIMs.

### 3D modeling of the bacterial chromosome structures

All the above results were based on chromatin interaction frequencies, what would they be if using real 3D structure models of chromosomes? To answer this, the 3D structure models of *E. coli* and *B. subtilis* chromosomes were constructed by using the EVR software, which is a 3D modeling tool particularly developed for bacterial chromosomes and has high accuracy and robustness [31]. Based on the same set of chromatin interaction data (**Figure S4**), the 3D chromosome structure models were constructed (**Figure 6**). The analysis for the global TRN (including positive and negative regulations) (**Figure 1**), the hierarchical structure of TRN (**Figure 2**) and the network motifs in TRN (**Figure 3–5**) were also conducted based on the spatial distances in the 3D chromosome models (**Figure S5-S9**). It could be found that, although not completely identical, the comparison results based on the 3D spatial distance are quite consistent with those based on the chromatin interaction frequency. What should be noticed is the inverse relationship between chromatin interaction frequency and the spatial distance, namely, the higher the interaction frequency, the shorter the spatial distance. This inverse relationship is reflected in the opposite trends in the corresponding figures.

**Figure 6.**
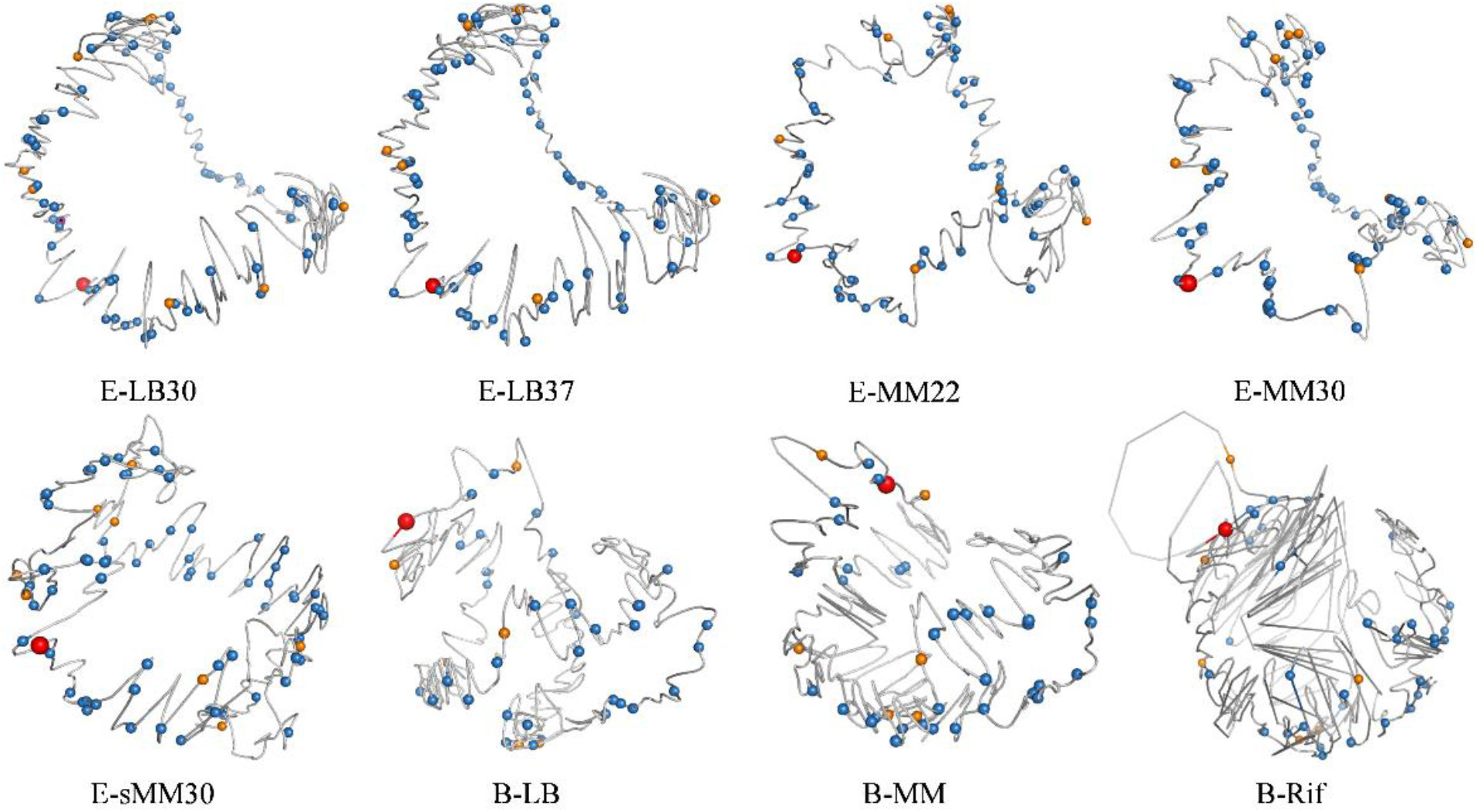
The distribution of high-level regulatory genes in the 3D architecture of bacterial chromosomes. In the 3D models of *E. coli* and *B. subtilis* chromosomes, the genes of the Top (orange) and Middle (blue) layers in the TRN hierarchy are shown as spheres, and the bigger red balls indicate the positions of oriC loci. The shape of *E. coli* chromosome looks like a letter C, and the top-layer genes (orange) are mainly distributed at both ends of the letter C; The shape of *B. subtilis* chromosome looks like a spiral, and the top-layer genes (orange) are mainly distributed in the middle and one end of the spiral. The middle-layer genes (blue) seem evenly distributed in the 3D architecture.

### Spatial effect in the dynamic function of network motifs

To illustrate the effect of spatial organization in gene regulation, we focused on network motifs and developed spatial models for their dynamic functions. By introducing the spatial positions of genes into the dynamic equations, we studied the effect of spatial distance between genes in network motifs. For FFL, the positions of three genes X, Y, Z can determine a plane, and the concentrations of the gene products (proteins) in the two-dimensional space are denoted as *X*_(*x*, *y*)_, *Y*_(*x*, *y*)_, *Z*(*x*, *y*), respectively. Without losing generality, we use a type-I coherent FFL (C1-FLL) [7] with AND gate to illustrate the spatial effect. For such a FLL, the gene Y can be activated by gene X when the concentration of the gene X product at the position of gene Y is higher than a threshold *δ_XY_*, and the gene Z can be activated by a combination of gene X and gene Y (namely, AND gate) when the concentrations of the products of gene X and gene Y at the position of gene Z are higher than the corresponding thresholds *δ_XZ_* and *δ_YZ_*, respectively. Considering the diffusion process, the product of a source gene (regulator) needs to spread to the target gene, and the further the distance between the two genes, the lower the concentration of the product of source gene when it reaches the position of the target gene. Therefore, the dynamic equations of such a “C1-FFL with AND gate” are as follows:

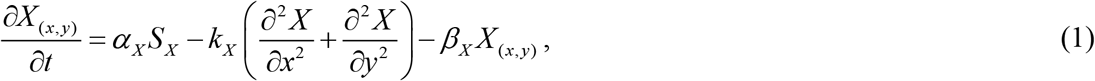

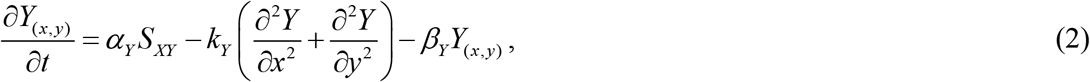

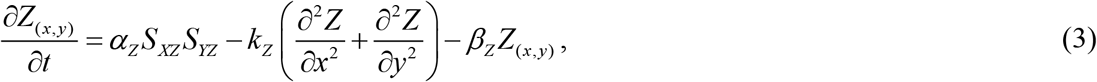

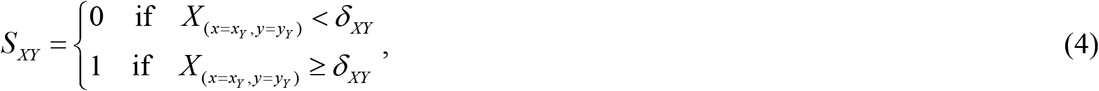

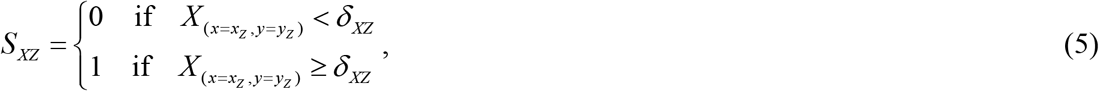

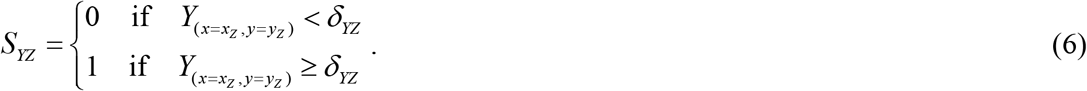

In this equation set, *α_X_/β_X_/k_X_*, *α_Y_/β_Y_/k_Y_*, *α_Z_/β_Z_/k_Z_* are the production/degradation/diffusion rates of the products of genes X, Y, Z, respectively, and *S_X_* is the signal to instantaneously transform the gene X product from inactive state to active state, and *S_XY_*, *S_XZ_*, *S_YZ_* are the activation signals for gene Y and gene Z which depend on the corresponding thresholds as listed in equations (4) – (6). Suppose the Euclidean distances between genes are denoted as *D*_X_Y_, *D*_X_Z_ and *D*_Y_Z_. To illustrate the influence of spatial distance on FFL function, we designed two FFLs with different spatial organization. As shown in **Figure 7A**, for FFL-1 (grey edges), suppose *D*_X_Y1_ = *D*_X_Z1_ = 2√2, *D*_Y1_Z1_ = 4; for FFL-2 (yellow edges), suppose Y2, Z2 are relatively far from X (*D*_X_Y2_ = 2√5, *D*_X_Z2_ = 3√2) and close to each other (*D*_Y2_Z2_ = √2). Except the spatial distribution, the other parameters for these two FFLs are identical. By solving the equation set (1) – (6) (see **Table S5** for parameter setting), the time course curves for the two FFLs are shown in **Figure 7A**. As can be seen, for the two pulse (*S_X_*) signals of shorter and longer duration, FFL-1 responds to both the two signals, while FFL-2 just responds to the longer one, which means the longer spatial distances between X_Y2 and X_Z2 in FFL-2 help it filter the signal but meanwhile the shorter spatial distance between Y2 and Z2 can accelerate the response of Z2.

**Figure 7.**
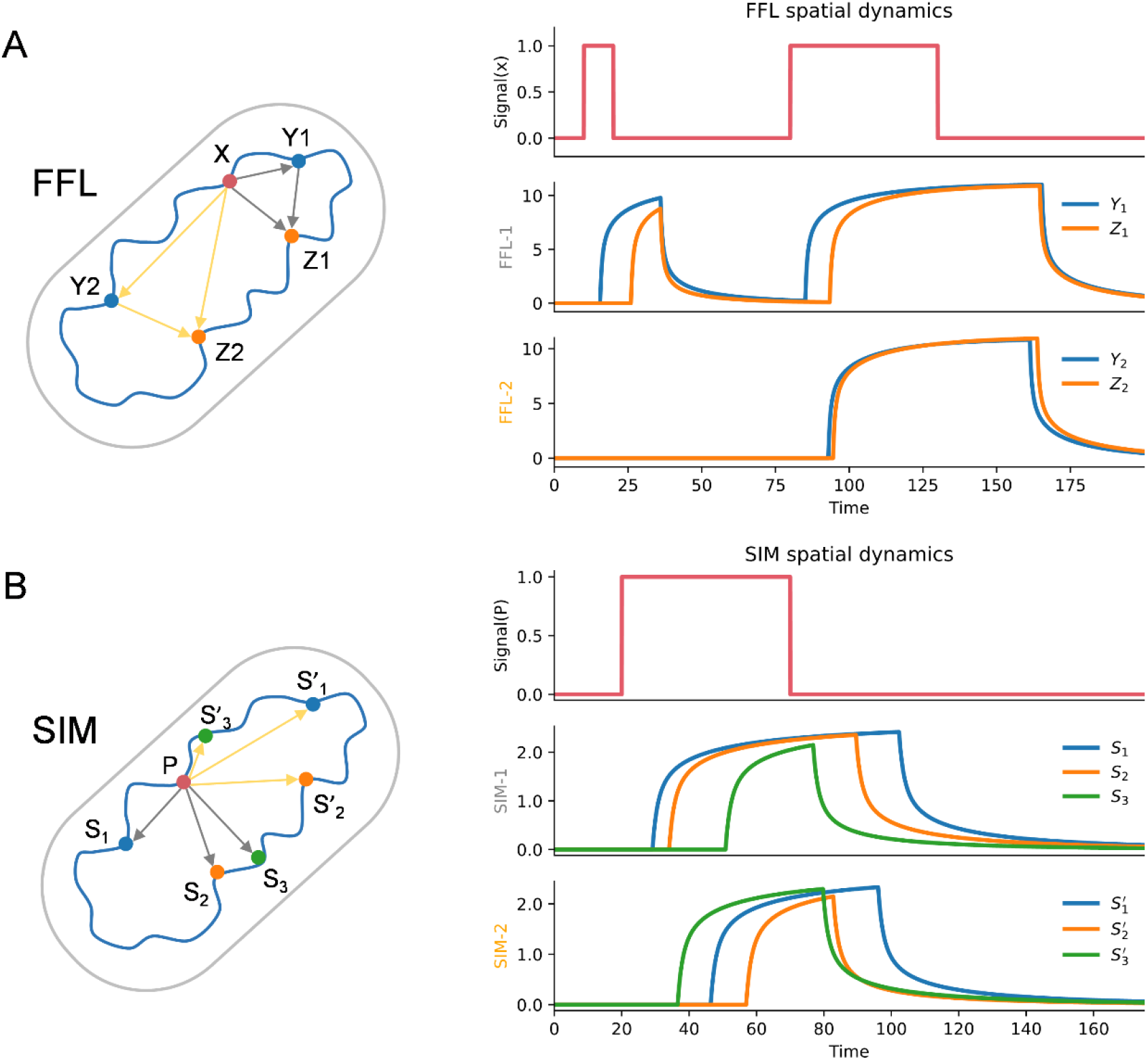
Dynamic behaviors affected by the spatial organization of FFL and SIM. (A). For FFL-1 (grey edges), suppose *D*_X_Y1_ = *D*_X_Z1_ = 2√2, *D*_Y1_Z1_ = 4; for FFL-2 (yellow edges), suppose *D*_X_Y2_ = 2√5, *D*_X_Z2_ = 3√2, *D*_Y2_Z2_ = √2 (see **Table S5** for the coordinates of X, Y1, Z1, Y2, Y2). FFL-1 responds to both the two signals, while FFL-2 just responds to the longer one, which means different spatial distribution of genes in FFL can affect dynamic function. (B). In SIM-1 (grey edges), the distances between the three child nodes (S_1_, S_2_, S_3_) and the parent node P are assumed to be identical, and the three child nodes can be regulated in a correct time order. In SIM-2 (yellow edges), the distances between the child nodes and the parent node are randomly arranged, and the original time order disappears. This result indicates that the spatial distribution of genes in SIM can affect dynamic function.

Similarly, for SIM with P (parent) as the regulator and S (son) as the target gene, we have the following dynamic equations:

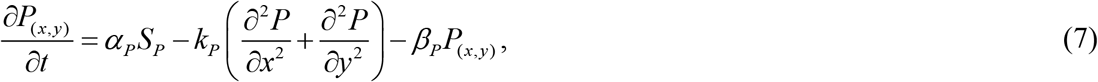

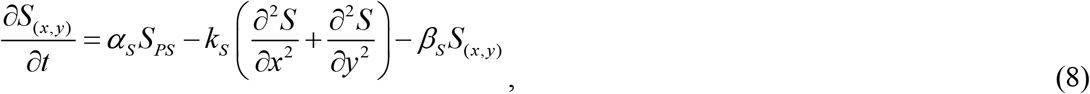

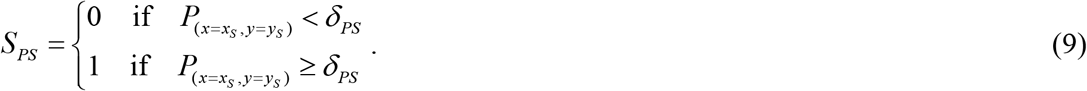

The parameters in this equation set have the same meaning as the corresponding parameters in FFL except that the symbols X, Y are replaced with the symbols of P and S. SIM has the function of sequentially switching ON/OFF the target genes in a temporal order. To illustrate the effect of spatial distance on this function, we designed two SIMs with different spatial distribution. As shown in **Figure 7B**, for SIM-1 (grey edges), the spatial distances between the regulator P and the targets *S_i_* (*i* = 1, 2, 3) are assumed to be identical (*D*_P_S1_ = *D*_P_S2_ = *D*_P_S3_), while for SIM-2 (yellow edges), the spatial distances are assumed to be all different (*D*_P_S1′_ > *D*_P_S2′_ > *D*_P_S3′_). Suppose the activation thresholds of the target genes are as follows: *δ*_P_S1_ = *δ*_P_S1′_ < *δ*_P_S2_ = *δ*_P_S2′_ < *δ*_P_S3_ = *δ*_P_S3′_. By solving the equation set (7) – (9) (see **Table S5** for parameter setting), the time course curves for the two SIMs are shown in **Figure 7B**. As can be seen, SIM-1 switches on the targets in a correct time order (namely, 1-2-3 sequentially), while the time order for SIM-2 is not correct, which means the uniformity of distances of the regulation edges is important for SIMs to function properly. The results of dynamic behavior analysis for FFL and SIM clearly indicate that the spatial distribution of genes in network motifs has great effects on their biological functions.

## Discussion

Previous studies on 3D genomes mainly focused on the organization and function of chromatin conformation [32]. Although it has been noticed that the spatial distance between a TF and its target gene may affect the efficiency and reliability of transcriptional regulation [9–12], these studies had to be concentrated on several or a small set of genes due to the lack of genome-wide information for gene spatial position. This work for the first time combined chromatin interaction data with the TRNs of *E. coli* and *B. subtilis* and disclosed the spatial organization features of the bacterial TRNs, which provided new insights into prokaryotic 3D genomics and systems biology.

The result that chromatin interaction between genes with regulation relationship is stronger than that of the genomic overall is in line with our previous finding that chromatin interaction within biological pathways is not random [14] and supports the connection between 3D genome organization and gene regulation [33]. It was found that positive regulation requires a larger but negative regulation requires a smaller spatial distance compared with the TRN, which indicates different regulation types have different requirements on the spatial arrangement of genes in 3D space to better fulfill their functions. It was also found that the chromatin interactions within FFLs are even stronger than the chromatin interaction of TRN and the chromatin interaction on the Y to Z edge is stronger than that on the other two edges of a FFL, which is quite stable among different experimental conditions. Previous work has shown that FFLs especially coherent FFLs can enhance the robustness of TRN [34], coupled with our findings in this work, indicating that FFLs are important for the organization and maintenance of bacterial TRNs. Although the chromatin interaction frequency within SIM is of no significant difference from that of TRN, it has higher uniformity, which may be important for a SIM to function as a timer with least interference from the spatial distance differences between the regulator and all its target genes. The spatial models of FFL and SIM dynamics were developed to rationalize the above findings (explanations) and the simulation results clearly demonstrate the effects of spatial distance (namely, the diffusion process of regulator) on the functions of these network motifs.

Our results are based on the chromatin interaction data and the TRNs of two bacteria species. Similar analysis may be extended to other prokaryotes and eukaryotes when data are available, to examine if these features are general among organisms. Actually, our previous work in yeast has found opposite overall trend as compared with the current bacterial results [35], possibly due to the obvious difference between eukaryotes and prokaryotes (namely, the existence of karyotheca), but more species should be studied to obtain general conclusion. Meanwhile, our findings suggest that the spatial arrangement of genes has important effects on their regulation relationships and thus their biological functions, and the spatial effect may be exploited in practice to modulate gene expression based on 3D distance through gene editing approaches such as TALEN or CRISPR [36, 37], which lays a foundation for the spatial-distance-based gene circuit design in synthetic biology [38].

## Materials and methods

### Chromatin interaction data

The chromatin interaction data of *E. coli* were derived from the original interaction matrix of 3C-seq for wild type MG1655 strain in the work by Lioy *et al*. [22]; the dataset corresponds to 5 culture conditions: LB medium early exponential phase at 30°C and 37°C (E-LB30, E-LB37), minimal medium early exponential phase (OD_600_ = 0.2) at 22°C and 30°C (E-MM22, E-MM30) and minimal medium stationary phase (OD_600_ = 2) at 30°C (E-sMM30). The chromatin interaction data of *B. subtilis* were derived from the Hi-C dataset of wild-type *Bacillus subtilis* HM1320 in the work by Marbouty *et al*. [23]; the dataset corresponds to 3 culture conditions: LB medium (B-LB), minimal medium (B-MM) and LB medium cultured then rifampicin treated for 20 minutes (B-Rif); all cells were grown at 30°C and harvested during the exponential phase (OD_600_ = 0.3). The original chromatin interaction matrixes of *E. coli* and *B. subtilis* were normalized using the Sequential Component Normalization (SCN) method [39]. The normalized interaction matrixes are shown in **Figure S4**.

### Definition of bins and localization of genes

The spatial distance of two genes is represented by the chromatin interaction frequency between two bins (a bin is defined as a DNA fragment of fixed length) that the genes belong to. The bin size is 5Kb for *E. coli* and 4Kb for *B. subtilis*, respectively. When the gene length is less than the length of a single bin, if a gene has more than a half of its total length in a bin, then the gene belongs to that bin; when the gene length is greater than the length of a single bin, if the gene length in a bin exceeds half of the bin length, then the gene is considered to contain that bin. For the interaction frequency between two genes spanning multiple bins, the average interaction frequency over all the bins the two genes involve is taken as the interaction frequency between the two genes. In data comparison, the statistical significance test was performed using the function wilcox.test in R package. The following convention symbols indicating significance levels were used: ns: *p* > 0.05; *: *p* <= 0.05; **: *p* <= 0.01; ***: *p* < = 0.001; ****: *p* <= 0.0001. The default significance level is 0.05 unless otherwise specified.

### Constructing 3D models of bacterial chromosomes

The normalized interaction matrixes were input into the EVR software [31] for the reconstruction of 3D chromosome structure models, and the default parameters were adopted: smoothing factor = 2, alpha = 0.5, itemsize = 20.0, mis_dis = 0.001, max_dis = 0.2, iter_num = “auto”, number of threads = “auto”, seed = “auto”. The generated 3D structure models of bacterial chromosomes were visualized using the PyMol software, and the visualization results are shown in **Figure 6**. The spatial distance between genes is defined as the Euclidean distance calculated based on the 3D coordinates of genes (bins) in space.

### Construction of transcriptional regulatory networks

The TRN of *E. coli* was constructed based on the transcriptional regulation experimental dataset from RegulonDB (version 9.0) [27]. After combining all regulatory relationships, the resulting TRN contains 2273 protein-coding genes (207 transcription factors, 7 sigma factors) and 6788 regulatory edges. The transcriptional regulation data of *B. subtilis* was adopted from SubtiWiki 2.0 [28], and the extracted gene type was protein-coding or pseudogenes. The resulting TRN contains 2400 nodes (including 192 transcription factors) and 5028 regulatory edges. Since chromatin interaction is what we concern, a regulatory edge was abandoned if its two nodes (regulator and target) are located in the same bin of chromatin partition. Consequently, there are 4562 positive (activation) and 1959 negative (repression) regulatory edges in the *E. coli* TRN; there are 3255 positive and 1714 negative regulatory edges in the *B. subtilis* TRN.

### Determination of network hierarchy

This study used the method from a previous publication [40] to divide the bacterial TRNs into four layers: Top, Middle, Bottom, and Target. The division algorithm is as follows. (1) The genes with no out-degree in the TRN are classified as the Target layer (TG), which are mainly the genes encoding enzymes and structural components of the cell; removal of all genes in the Target layer resulted in a TRN containing only transcription factors. (2) In the resulted TRN of only transcription factors, the gene set with “strongly connected component (SCC)” attribute is collapsed into a single super node, where a SCC is a fully connected subnetwork in which each pair of nodes u and v must have u to v and v to u edges; self-regulation loop is also a kind of SCC. The edges of other nodes in the network that are connected to the genes in SCCs are replaced by the connections of the former to the super nodes, thereby resulting into a directed acyclic graph. (3) In the directed acyclic graph, the nodes with no in-degrees are classified as the Top layer (T); the nodes with no out-degrees are classified as the Bottom layer (B), and the isolated transcription factors are also classified into this layer; the remaining nodes are classified into the Middle layer (M).

### Discovery of network motifs

The software mfinder1.21 [41] was used in searching FFLs in the *E. coli* and *B. subtilis* TRNs with parameter settings: r = 1000 times. In the search results, FFLs were identified as network motifs with occurrence frequency = 4599 and z-score = 4.88 for *E. coli* and occurrence frequency = 2066 and z-score = 9.50 for *B. subtilis*. SIM was defined according to the publication by Konagurthu and Lesk [42]: only one parent (regulating) node and at least two children (regulated) nodes and the in-degree (number of incoming edges) for each child node strictly equal one within the full network not just within the sub-graph. According to this definition, 30 and 58 SIMs were identified in the *E. coli* and *B. subtilis* TRNs, respectively.

## Abbreviations

TRN: transcriptional regulatory network
TF: transcription factor
FFL: feed-forward loop
SIM: single input module
SD: standard deviation
3D: three-dimensional

## Acknowledgements

This work was supported by the National Natural Science Foundation of China (Grant 31971184). The funders had no role in study design, data collection and interpretation, or the decision to submit the work for publication. We thank Hong-Rui Xu for valuable information.

## Author Contributions

Conceived and designed the research: B.G.M. Performed the experiments: L.T., T.L., K.J.H., X.P.H. Analyzed the data: L.T., T.L, K.J.H, B.G.M. Wrote the manuscript: L.T., B.G.M. Revised the manuscript: X.P.H.

## Additional Information

**Supplementary information** accompanies this paper at http://

