## Supplementary Figures for "The Spatial Organization of Bacterial Transcriptional Regulatory Networks"

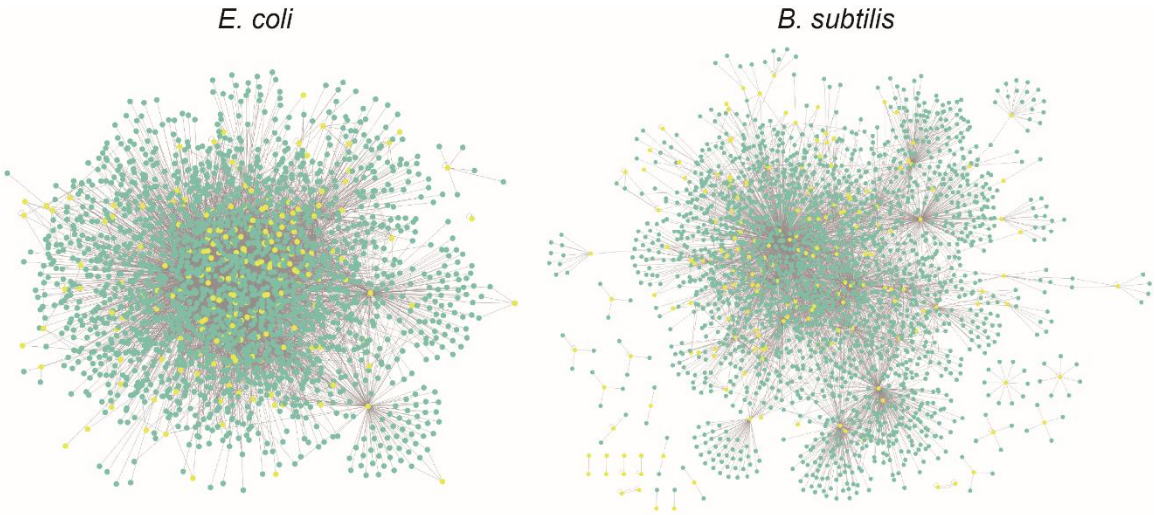


**Figure S1.** The reconstructed transcriptional regulatory networks (TRNs) of *Escherichia coli* and *Bacillus subtilis.* Nodes of yellow color are transcription factors.

| Key Nodes | E-LB30 | E-LB37 | E-MM22 | E-MM30 | E-sMM30 | B-LB | B-MM | B-Rif |
|---|---|---|---|---|---|---|---|---|
| In-Hub-I | > 1.74E-09 | > 1.55E-05 | > 6.88E-09 | > 9.23E-07 | > 5.95E-08 | > 7.05E-06 | > 0.0207 | = 0.1293 |
| In-Hub-O | = 0.9408 | = 0.9676 | < 1.93E-05 | < 0.0003 | < 0.0229 | = 0.4985 | = 0.2325 | < 0.0057 |
| Out-Hub-I | > 0.0450 | > 0.0392 | = 0.1414 | = 0.1333 | = 0.1968 | > 0.0001 | > 0.0005 | > 0.0020 |
| Out-Hub-O | < 1.73E-15 | < 1.07E-16 | < 2.27E-15 | < 8.27E-15 | < 1.02E-14 | < 3.30E-17 | < 2.90E-17 | < 1.87E-14 |
| Bottleneck-I | > 0.0016 | > 0.0040 | > 0.0132 | > 0.0155 | > 0.0095 | > 0.0008 | > 0.0003 | > 4.38E-05 |
| Bottleneck-O | < 2.25E-14 | < 4.86E-15 | < 2.83E-14 | < 1.21E-13 | < 4.46E-14 | < 3.46E-12 | < 1.97E-14 | < 4.03E-12 |
| Center-I | > 0.0040 | = 0.3958 | = 0.5635 | = 0.1122 | > 0.0039 | NA | NA | NA |
| Center-O | > 0.0002 | > 0.0420 | = 0.4143 | = 0.6267 | > 0.0063 | < 0.0067 | = 0.2043 | = 0.5624 |

**Figure S2.** Chromatin interaction frequencies of key nodes compared with TRN by using Wilcoxon rank sum tests. "-I" and "-O" after In-Hub, Out-Hub, Bottleneck, Center represent the chromatin interaction "intra" the 4 kinds of key nodes and the chromatin interaction between the 4 kinds of key nodes and "other" genes, respectively. ">", "<", "=" represent the chromatin interaction frequency is higher, lower, of no significant difference compared with that of TRN, respectively; the number below them are *p*-values in Wilcoxon rank sum tests for statistical significance. Red: significantly higher than TRN; Green: significantly lower than TRN.

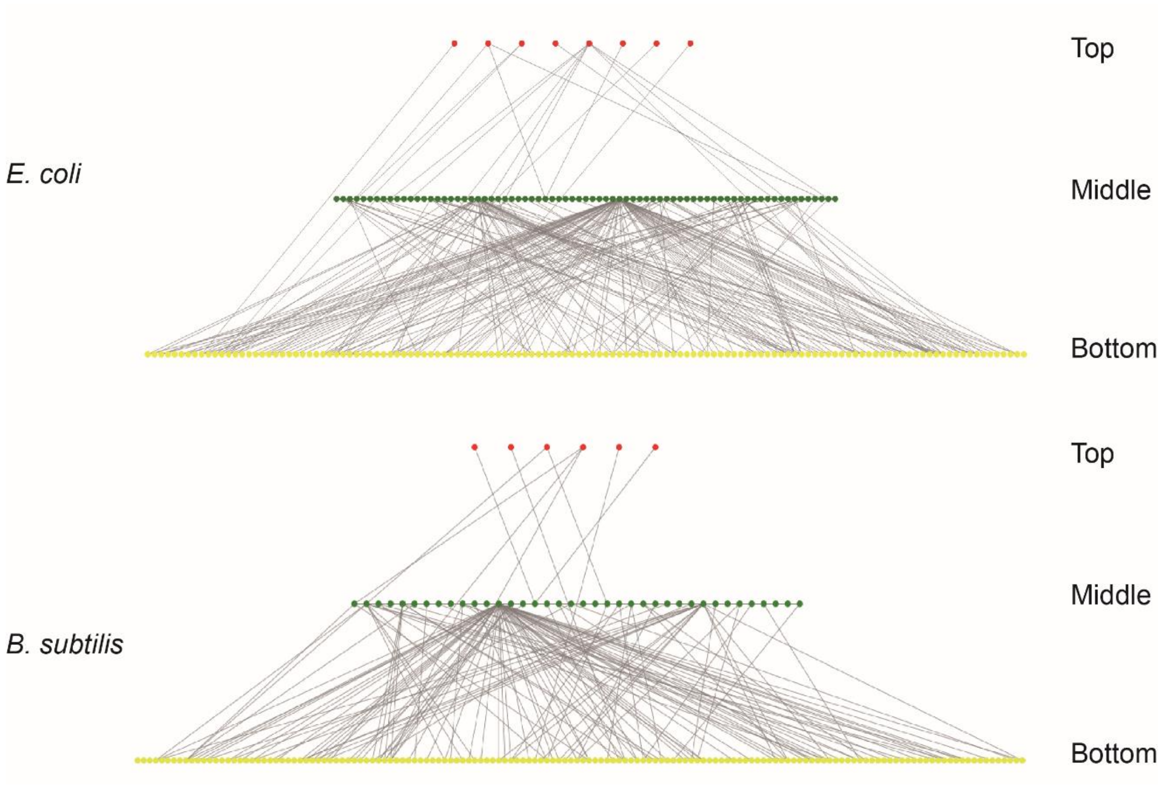


**Figure S3**. The hierarchical structures of *Escherichia coli* and *Bacillus subtilis* TRNs.

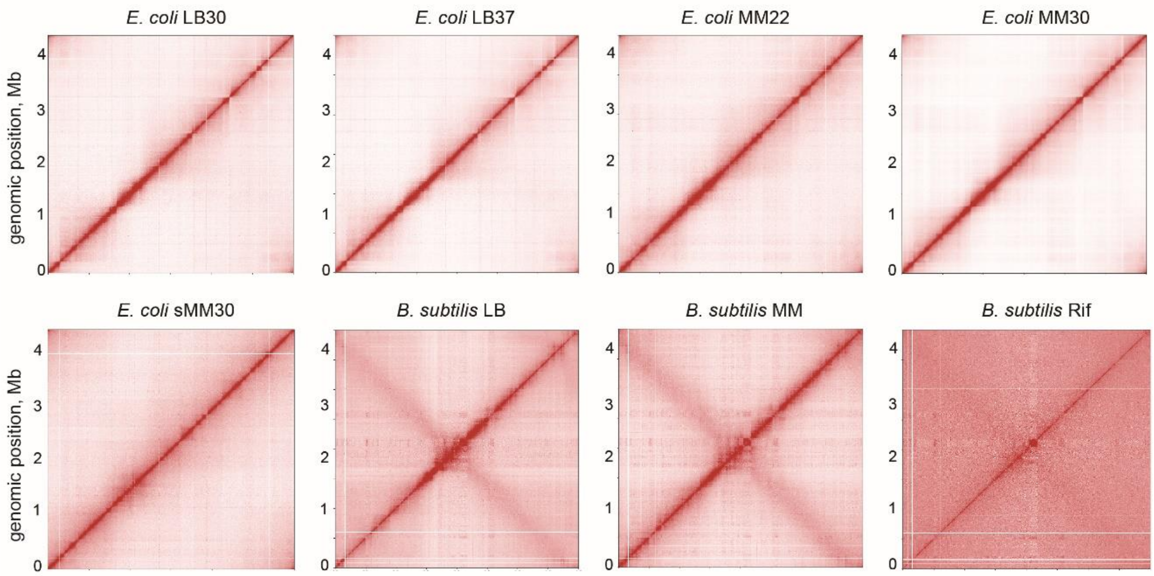


**Figure S4.** The normalized chromatin interaction frequency matrixes used in this study.

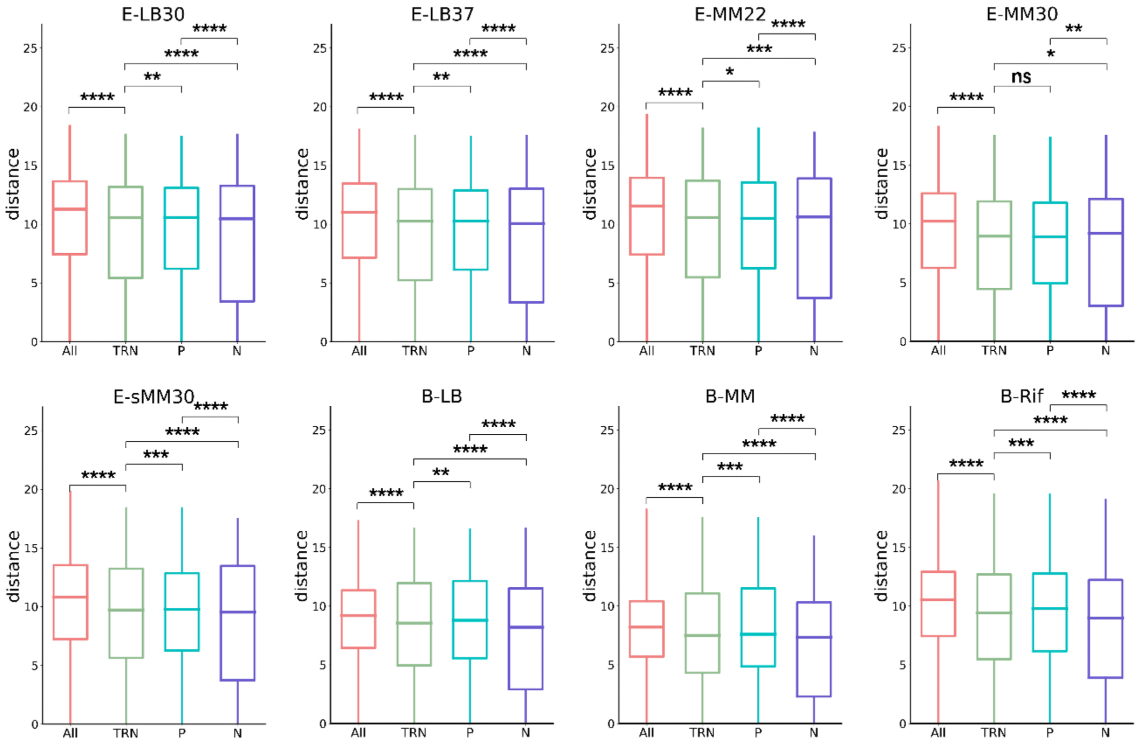


**Figure S5. Comparison of spatial distances between gene pairs in global TRN.**

In almost all culture conditions, the spatial distances within TRN (denoted as TRN) are significantly shorter than those between all the gene pairs in the whole genome (denoted as All); the spatial distance of positive regulation (P) is significantly longer than TRN, while the spatial distance of negative regulation (N) is significantly shorter than TRN.

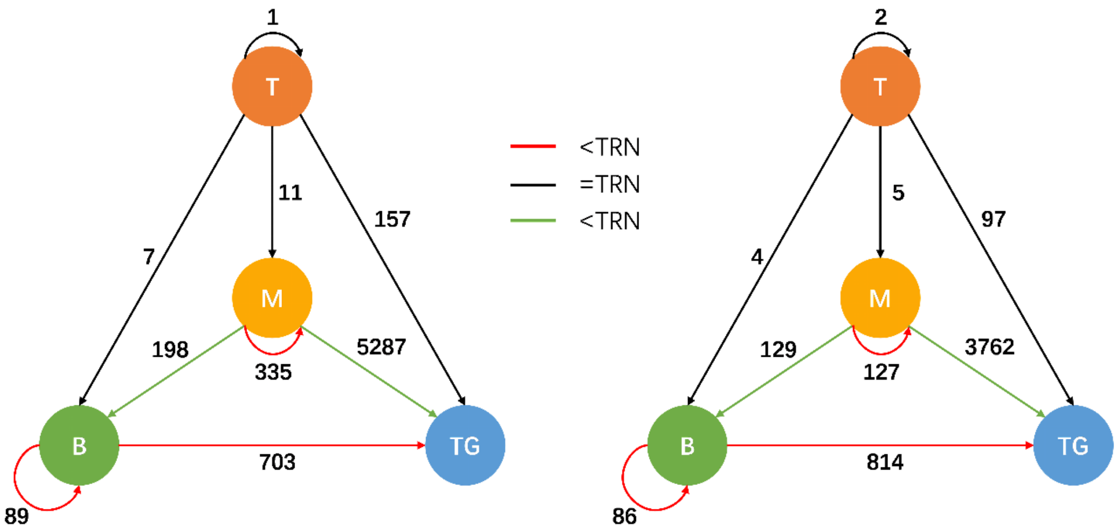


**Figure S6. The spatial organization of TRN hierarchy based on 3D distance.**

The four nodes T, M, B and TG represent the Top, Middle, Bottom and Target layers in the hierarchy, respectively. The numbers on the edges represent the numbers of regulatory relationships within or between layers. The color of edge represents the result of comparing the spatial distance between the gene pairs of the edge with TRN. Significance level: $p < 0.05$.

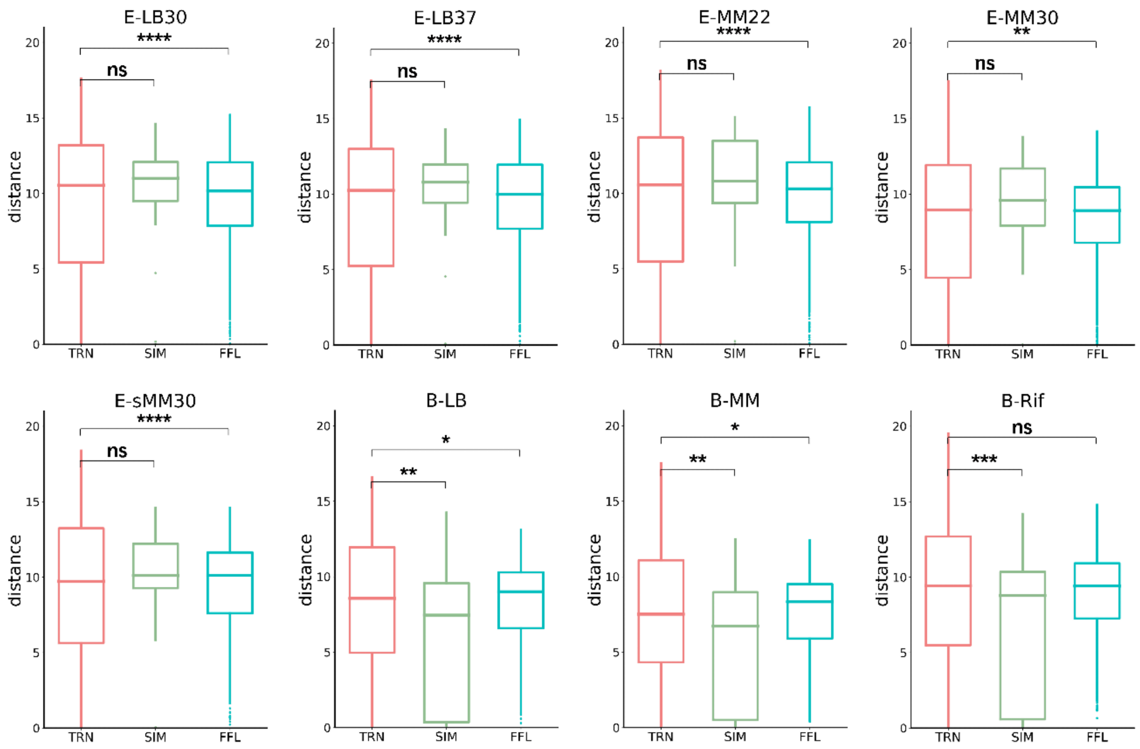


**Figure S7. Comparison of spatial distances between gene pairs in network motifs with TRN.**

In the five culture conditions of *E. coli*, the spatial distance of FFL is significantly shorter than that of TRN and the spatial distance of SIM is of no significant difference from TRN. For most cases in *B. subtilis*, the spatial distances between gene pairs of both kinds of network motifs are significantly shorter than that of TRN.

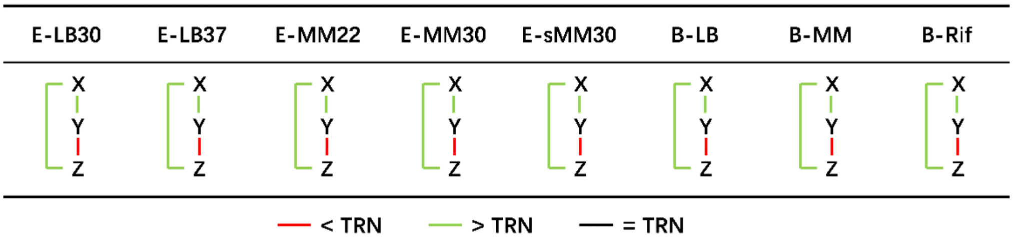


**Figure S8. Comparison of spatial distances between gene pairs in FFL edges with TRN.**

In all cases, the spatial distance between X and Y/Z in feed-forward loop is significantly longer than TRN, while the spatial distance between Y and Z is significantly shorter than TRN. Significance level: $p < 0.05$.

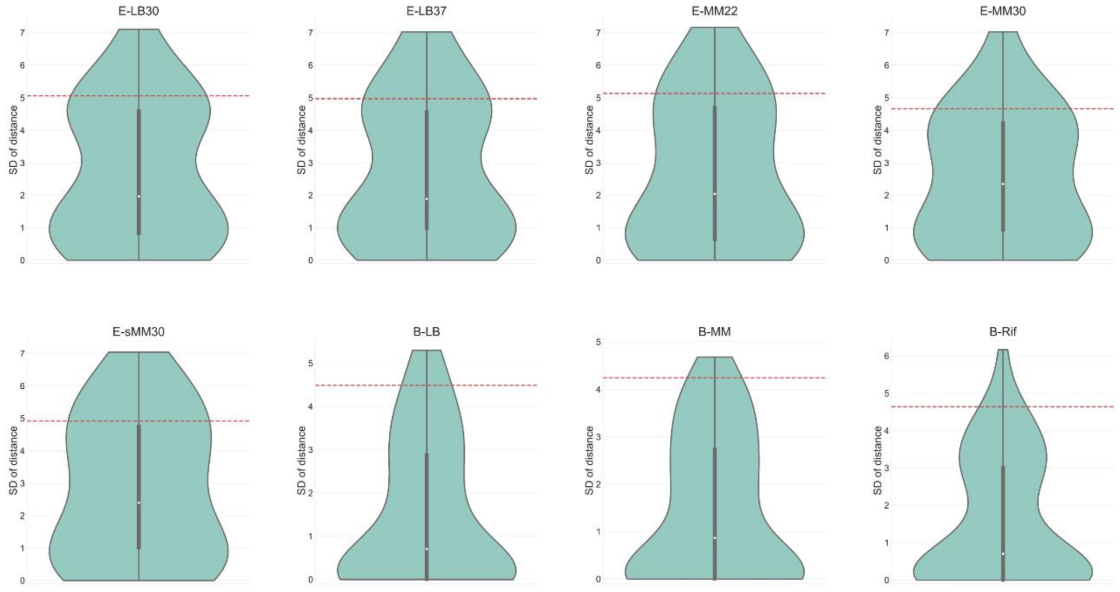


**Figure S9. The distribution of standard deviation (SD) of spatial distance within SIMs (violin plot) and its comparison with that of TRN (red dashed line).**

The SD of spatial distance within SIMs is apparently lower than that of TRN, indicating lower dispersion and higher uniformity of spatial distance within SIMs.
